# Intrathecal administration of palmitoyl-lysophosphatidylethanolamine reduces secondary injury and improves locomotor recovery following spinal cord injury

**DOI:** 10.1101/2025.09.01.673598

**Authors:** Tetsuhiko Mimura, Yusuke Tanikawa, Shiori Kawase, Taishi Kotani, Eiko Kato, Taiga Kurihara, Yoshikazu Matsuda, Naoto Saito, Jun Takahashi, Takeshi Uemura

**Author notes:** Correspondence (T.U.).

## Abstract

Spinal cord injury (SCI) triggers secondary pathophysiological cascades, including glutamate excitotoxicity, that result in neuronal loss and impair functional recovery. We have previously shown that lysophosphatidylethanolamine (LPE), a lysophospholipid, promotes neurite outgrowth and protects against glutamate excitotoxicity in cultured cortical neurons. However, whether these effects extend to spinal cord neurons and occur in vivo has remained unclear. In this study, we compared the effects of different LPE species: myristoyl-LPE (14:0 LPE), palmitoyl-LPE (16:0 LPE), stearoyl-LPE (18:0 LPE), and oleoyl-LPE (18:1 LPE) in cultured spinal commissural neurons, and evaluated their effects in vivo using a mouse model of SCI. In cultured neurons, all LPE species promoted neurite outgrowth. Although several species demonstrated a tendency toward neuroprotection, only 16:0 LPE exhibited a statistically significant protective effect against glutamate-induced excitotoxic cell death. Intrathecal administration of 16:0 LPE after SCI reduced TUNEL-positive cells in the acute phase and attenuated lesion expansion at 8 weeks post-injury. Moreover, 5-HT fluorescence intensity was increased in 16:0 LPE-treated mice, suggesting enhanced serotonergic innervation. Furthermore, administration of 16:0 LPE after SCI significantly improved hind-limb motor performance compared with vehicle controls, as assessed by the Basso Mouse Scale. Collectively, these findings suggest that intrathecal administration of 16:0 LPE reduces secondary injury and promotes functional recovery following SCI. Our findings highlight its potential as a therapeutic candidate for SCI.

## Introduction

Spinal cord injury (SCI) results from fractures, dislocation, distraction, or compression of the vertebral column that cause damage to the spinal cord. Although contusion–compression injuries may produce only partial deficits, SCI frequently leads to severe and permanent impairments in motor, sensory, and autonomic functions. Globally, more than one million individuals are estimated to live with paresis due to SCI (van den Berg et al., 2010).

Traumatic SCI proceeds through two distinct phases: primary and secondary injury. The primary phase consists of irreversible mechanical damage, including fracture or dislocation, that causes neurological dysfunction at and below the site of impact. This initial insult initiates a cascade of secondary events, including ischemia, disruption of the blood–spinal cord barrier, neuroinflammation, oxidative stress, and glutamate-mediated excitotoxicity (Tator and Fehlings, 1991; Wilson et al., 2013; Nagoshi and Okano, 2018). These processes interact and collectively exacerbate tissue damage, resulting in progressive axonal degeneration and neuronal loss. Various therapeutic approaches have been investigated to mitigate secondary injury following SCI (Ashammakhi et al., 2019). However, a universally effective treatment has not yet been identified (Zeb et al., 2016).

Phospholipids are amphiphilic molecules that play a critical roles in membrane architecture and intracellular signaling (van Meer et al., 2008). Among these, lysophospholipids have been identified as significant bioactive mediators (Makide et al., 2009). Lysophosphatidylethanolamine (LPE), one of the lysophospholipids, generated from phosphatidylethanolamine by phospholipase A–type reactions, has been detected in human plasma and cerebrospinal fluid at submicromolar to micromolar concentrations (Liu et al., 2015; Sato et al., 2010; Péter et al., 2020; Yamamoto et al., 2021). Changes in LPE levels have been observed in rodent models of traumatic brain injury and cognitive dysfunction (Guo et al., 2017; Villamil-Ortiz et al., 2016; Sabogal-Guáqueta et al., 2018), suggesting potential physiological and pathological relevance in the central nervous system.

Our previous studies demonstrated that 16:0 LPE, 18:0 LPE, and 18:1 LPE promote neurite outgrowth in cultured cortical neurons (Hisano et al., 2021a). Additionally, 18:1 LPE protects against glutamate-induced toxicity in these cultures (Hisano et al., 2021b). However, it remains unclear whether these LPE species exert similar effects in spinal cord neurons. We hypothesized that, if LPEs exhibit such effects in spinal cord neurons, they may mitigate excitotoxic injury and thereby improve functional outcomes following SCI.

In the present study, we examined the effects of different LPE species in cultured spinal commissural neurons and identified palmitoyl-LPE (16:0 LPE) as the most effective. We further investigated the in vivo effects of intrathecal administration of 16:0 LPE in a mouse contusion model of SCI. Our findings indicate that 16:0 LPE reduces neuronal apoptosis in the acute phase, attenuates chronic lesion expansion, enhances serotonergic innervation, and improves locomotor recovery. These results support 16:0 LPE as a candidate therapeutic agent for SCI and extend our understanding of its functions in the central nervous system.

## Materials and methods

### Animal Care and Use

The animal experimental protocols were reviewed by the Committee for Animal Experiments and approved by the President of Shinshu University (Approval Nos. 021019 and 023123), and all experiments were conducted in accordance with the Guidelines for the Care and Use of Laboratory Animals of Shinshu University.

### Cell Cultures

Spinal commissural neurons were prepared from ICR mouse embryos at embryonic day 12 (E12) as previously described (Langlois et al., 2010). Pregnant ICR mice were purchased from Japan SLC Inc. (Shizuoka, Japan). Briefly, E12 embryos were dissected from the uterus into a 10-cm Petri dish containing ice-cold Leibovitz’s L-15 medium (FUJIFILM Wako Chemicals). Spinal cords were isolated using fine forceps and carefully dissected free of dorsal root ganglia. The isolated spinal cord was arranged in an “open book” configuration within a Petri dish containing Leibovitz’s L-15 medium supplemented with 10% heat-inactivated horse serum (HiHS). The spinal cord was then secured in place using fine forceps. Subsequently, an L-shaped tungsten needle was employed to make an incision in the dorsal region, excising approximately one-fifth of the width of one side of the spinal cord. The excised dorsal tissue was then transferred to a 15 mL tube containing Leibovitz’s L-15 medium with 10% HiHS and maintained on ice until further processing. Following a wash with Hank’s Balanced Salt Solution (HBSS), tissues were incubated in 1% trypsin (Sigma-Aldrich) and 0.1% DNase I (SIGMA) in Ca^2+^/Mg^2+^-free phosphate buffered saline (PBS) for 5 min at room temperature. The cells were washed with a culture medium containing 5% fetal bovine serum (FBS; Gibco) and dissociated by passing through a fire-polished Pasteur pipette in Ca^2+^/Mg^2+^-free PBS containing 0.05% DNase I, 0.03% trypsin inhibitor (Sigma-Aldrich), and 2 mM MgCl_2_.

Dissociated cells were collected by low-speed centrifugation and resuspended in plating medium consisting of Neurobasal medium (Thermo Fisher Scientific) supplemented with 5% fetal bovine serum (FBS), 100 U/ml penicillin, 100 μg/ml streptomycin, and 0.2 mM GlutaMAX-I (Thermo Fisher Scientific). Sixteen hours after plating, the medium was replaced with Neurobasal medium supplemented with 2% B-27 (Thermo Fisher Scientific), 100 U/ml penicillin, 100 μg/ml streptomycin, and 0.2 mM GlutaMAX-I. All subsequent culture steps were carried out under serum-free conditions. For morphological analyses, cells were plated at a density of 2.0 × 10^4^ cells/well on 12-mm diameter coverslips precoated with 30 μg/ml poly-L-lysine (70–150 kDa; Sigma-Aldrich) in 24-well culture plates. For glutamate toxicity assays, cells were plated at a density of 7.8 × 10^4^ cells/well in 24-well plates. To inhibit glial proliferation, 4 μM cytosine β-D-arabinofuranoside (Ara-C) was added to the culture medium at days in vitro (DIV) 3. At DIV6, half of the culture medium was replaced with fresh serum-free medium.

### Morphological Analysis of Cultured Neurons

To examine the effects of phospholipids on neurite outgrowth, primary neuronal cultures were treated at DIV1 with 1 μM of the following LPEs: 14:0 LPE (1-myristoyl-2-hydroxy-sn-glycero-3-phosphoethanolamine; Cayman Chemical), 16:0 LPE (1-palmitoyl-2-hydroxy-sn-glycero-3-phosphoethanolamine; Echelon Biosciences), 18:0 LPE (1-stearoyl-2-hydroxy-sn-glycero-3-phosphoethanolamine; Avanti Polar Lipids), and 18:1 LPE (1-oleoyl-2-hydroxy-sn-glycero-3-phosphoethanolamine; Cayman Chemical). Cultures were fixed at DIV2, DIV3, and DIV4 with 4% paraformaldehyde (PFA) and 4% sucrose in PBS. After fixation, cells were permeabilized with PBS containing 0.25% Triton X-100 for 5 min and blocked with 10% donkey serum. The cultures were then immunostained with mouse anti-Tuj1/βIII-tubulin antibody (1:5,000; BioLegend), followed by Alexa Fluor 488-conjugated donkey anti-mouse IgG secondary antibody (1:500; Thermo Fisher Scientific).

### Glutamate Toxicity Assay

At DIV15, spinal cord cultured neurons were preincubated with 10 μM of 14:0 LPE, 16:0 LPE, 18:0 LPE, or 18:1 LPE for 1 h. The culture medium was then replaced with Locke’s buffer (NaCl, 154 mM; KCl, 5.6 mM; CaCl_2_, 2.3 mM; MgCl_2_, 1.0 mM; NaHCO_3_, 3.6 mM; glucose, 5 mM; HEPES, 5 mM; pH 7.2)(Du et al., 2007), containing 5 μM glutamate, and cells were incubated for an additional 1 h at 37°C. After glutamate treatment, the Locke’s buffer was removed, and the previously collected culture medium containing the respective LPEs was returned to the wells. After 24 h, the living cells were stained with calcein-AM (Dojindo Laboratories) according to the manufacturer’s instructions. For immunocytochemistry, cells were fixed and incubated with anti-NeuN antibody (1:500, Millipore), followed by incubation with Alexa Fluor 555-conjugated donkey anti-mouse IgG antibody (1:500, Thermo Fisher Scientific).

### SCI Model

Female C57BL/6J mice (8 weeks old, 17–20 g; Japan SLC, Hamamatsu, Japan) were used for SCI experiments. Mice were anesthetized with a mixture of 4 mg/kg midazolam, 0.3 mg/kg medetomidine, and 5 mg/kg butorphanol. Subsequently, complete laminectomy at the tenth thoracic vertebra (T10) vertebral level was performed. The dorsal surface of the dura mater was exposed, and a contusive SCI was induced at T10 using the Infinite Horizon Impactor (Precision Systems & Instrumentation) at a force of 70 kdyn, as previously described (Hara et al., 2017; Suzuki et al., 2020; Uezono et al., 2018). The strength and duration of the impact were monitored through the automated recording system integrated into the impactor. After injury, muscle and skin were closed in layers, and the mice were placed in a temperature-controlled chamber until thermoregulation was re-established. Postoperatively, mice were monitored daily for general health, mobility within the cage, wound healing, infection, and toe autophagy. Bladders were manually expressed twice daily for the first postoperative week and once daily until spontaneous voiding was restored.

### Intrathecal Administration of LPE

Mice subjected to SCI were randomly assigned to vehicle-treated or 16:0 LPE-treated groups. 16:0-LPE was dissolved in 0.2% DMSO in PBS at a final concentration of 10 µM. Vehicle-treated mice received an equivalent volume of vehicle. For intrathecal administration, a 26-gauge Hamilton syringe needle (Sigma-Aldrich) was inserted into the dural space through the interlaminar region between the fourth and fifth lumbar vertebrae, and 10 μl of solution (16:0 LPE or vehicle) was injected over 2 min. To minimize cerebrospinal fluid leakage, the needle was left in place at the injection site for an additional 2 min before withdrawal. Intrathecal injections were performed immediately after SCI and repeated at 1 and 2 days post-injury, for a total of 3 administrations. For terminal deoxynucleotidyl transferase-mediated dUTP nick end labeling (TUNEL) analysis, LPE was administered twice: immediately after injury and again on the following day, prior to tissue sample collection.

### Immunohistochemistry

After behavioral analysis 56 days after SCI, mice were anesthetized and perfused transcardially with 4% PFA in PBS. Spinal cords were dissected, post-fixed in the same fixative overnight, and subsequently immersed in 30% sucrose in PBS at 4°C for 2–3 days. Tissues were embedded in Tissue-Tek optimal cutting temperature (O.C.T.) compound (Sakura Finetek), and sectioned at a thickness of 20 μm using a cryostat (CM1900; Leica). For immunostaining, sections were incubated for 1 h at room temperature in blocking solution containing 10% normal donkey serum and 0.1% Triton X-100 in PBS. After blocking, the sections were incubated with rat anti-CD11b (1:500; clone 5C6, Bio-Rad), rabbit anti-glial fibrillary acidic protein (GFAP) (1:1000; Hisano et al., 2009) or rabbit anti-5-hydroxytryptamine (5-HT) (1:200; ImmunoStar) antibody, followed by incubation with Alexa Fluor 488-conjugated donkey anti-rabbit IgG (1:500; Thermo Fisher Scientific), Alexa Fluor 555-conjugated donkey anti-rabbit IgG (1:500; Thermo Fisher Scientific), or Cy3-conjugated goat anti-rat IgG (1:500; Jackson ImmunoResearch) antibody. TUNEL histochemistry was performed two days after SCI using the In Situ Cell Death Detection Kit, TMR Red (Roche), according to the manufacturer’s instructions. The sections were counterstained with 4′,6-diamidino-2-phenylindole (DAPI, Thermo Fisher Scientific).

### Image Acquisition and Quantitative Analysis

Images of culture experiments were taken with a confocal laser-scanning microscope (TCS SP8; Leica Microsystems) using either HC PL APO CS2 20×/0.75 NA multiple immersion lens or HC PL APO CS 10×/0.40 NA multiple immersion lens (Leica Microsystems). Images of immunohistochemistry were taken with a fluorescence microscope (BZ-X700; Keyence). A 10× objective lens (NA 0.45, Keyence) was used for immunostaining of GFAP, CD11b, and TUNEL, while a 20× objective lens (NA 0.45, Keyence) was used for immunostaining of 5-HT. Whole-slide images were generated by stitching multiple fields of view using Keyence BZ-X700. Exposure times and gain settings were kept constant across experimental groups. For lesion area quantification, five sagittal sections (20 μm thick, spaced 180 μm apart) were selected from each mouse, consisting of the section containing the lesion epicenter as well as two additional sections rostrally and two laterally on each side. The GFAP-negative volume surrounding the lesion was measured, and tissue volume was estimated using the Cavalieri principle(Akbas et al., 2004; Aleksić et al., 2014). GFAP and CD11b signals were evaluated using GFAP and CD11b signal intensity profiles within ±1.5 mm of the epicenter. For the analysis of serotonergic innervation, midsagittal sections were used to generate a rostro–caudal intensity profile of 5-HT immunoreactivity from 0.5 to 3.0 mm caudal to the epicenter. To evaluate apoptotic cell death, TUNEL-positive cells with DAPI-positive nuclei were counted within regions extending 2.5 mm rostral and 2.5 mm caudal to the epicenter. All signal intensities were background-subtracted prior to analysis. Background subtraction was performed using the “Subtract Background” plugin in Fiji (ImageJ, Version 1.54p) with a rolling-ball radius of 30 μm. All quantitative measurements were performed in ImageJ Fiji Version 1.54p.

### Behavioral Analysis

Locomotor recovery was evaluated from video recordings in the open field using the Basso Mouse Scale (BMS). Hindlimb motor function was scored weekly for 8 weeks post-injury. When right and left hindlimb scores differed, the BMS score was reported as the average of the two. All assessments were performed in a blinded manner with respect to the experimental groups.

### Statistical Analysis

Statistical significance was evaluated using two-way repeated-measures ANOVA followed by Welch’s t-test for post-hoc analysis or one-way ANOVA followed by Tukey’s HSD test. Two-group comparisons were analyzed using Welch’s t-test. Statistical analyses were performed using R software (version 4.2.1) (R Core Team, 2022) A *P* value < 0.05 was considered statistically significant.

## Results

### LPE species enhanced neurite outgrowth in cultured spinal commissural neurons

Our previous studies demonstrated that LPE species stimulate neurite outgrowth in cultured cortical neurons (Hisano et al., 2021a; 2021b). To examine whether LPE species exert similar effects in spinal commissural neurons, we treated cultured neurons at DIV1 with 1 μM 14:0, 16:0, 18:0, or 18:1 LPE, and fixed them for Tuj1 immunostaining at DIV2–DIV4 (Fig. 1A). In control culture, the length of neurites increased progressively from DIV2 to DIV4. Quantitative analysis of the longest neurite further revealed that all examined LPE species significantly enhanced neurite outgrowth (Fig. 1B), with no substantial differences observed among them. On the other hand, the number of neurites extending from the cell body was significantly increased compared to the control group only with the addition of 16:0 LPE and 18:1 LPE but not 14:0 LPE or 18:0 LPE. The total length of the neurites was significantly greater than the control in all LPE-treated groups. These results suggested that LPEs can promote the extension of neurites in cultured spinal commissural neurons.

**Figure 1.**
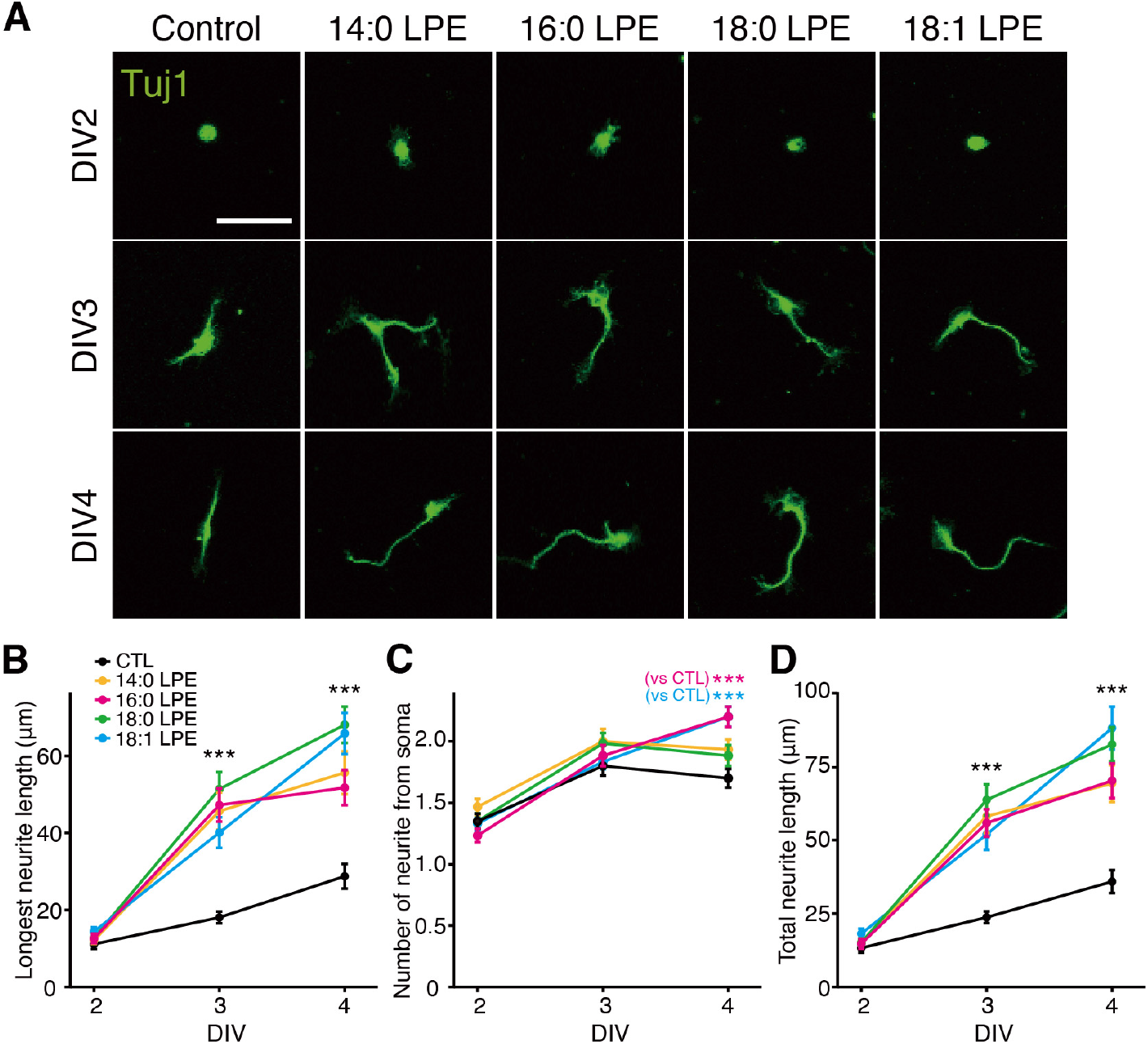
LPE species promote neurite outgrowth in cultured spinal commissural neurons. (A) Effects of various LPE species on neurite outgrowth in cultured spinal commissural neurons. Cultures were treated with 1 μM 14:0 LPE, 16:0 LPE, 18:0 LPE, or 18:1 LPE at DIV 1 and immunostained with an antibody against Tuj1 at DIV 2, 3, and 4. Representative images are shown. Scale bar, 50 µm. (B) Quantification of the length of the longest neurite emerging from the soma in (A). (C) Quantification of the number of neurites emerging from the soma in (A). (D) Quantification of the total neurite length of neurites in (A). All values represent mean ± SEM. n = 30 cells per group. ***p < 0.001; one-way ANOVA followed by post hoc Tukey’s test.

### 16:0 LPE protects cultured spinal commissural neurons from glutamate toxicity

Our previous work also showed that 18:1 LPE protects cultured cortical neurons against glutamate toxicity (Hisano et al., 2021b). To examine whether LPE species exert similar neuroprotective effects in spinal commissural neurons, we exposed cultures to 5 μM glutamate for 1 h and subsequently treated them with 10 μM 14:0, 16:0, 18:0, or 18:1 LPE (Fig. 2A). When the cultures were exposed to 5 μM glutamate for 1 h, the proportion of calcein/NeuN double-positive viable neurons was reduced to approximately 20% of that in control cultures after 24 h (Fig. 2B). Application of 16:0 LPE resulted in an approximate doubling of the number of viable neurons compared to cultures treated with glutamate alone. In contrast, 14:0 LPE, 18:0 LPE, and 18:1 LPE had little effect on the glutamate-induced reduction in viable neurons. These results suggested that 16:0 LPE protects spinal commissural neurons from glutamate-induced cell death.

**Figure 2.**
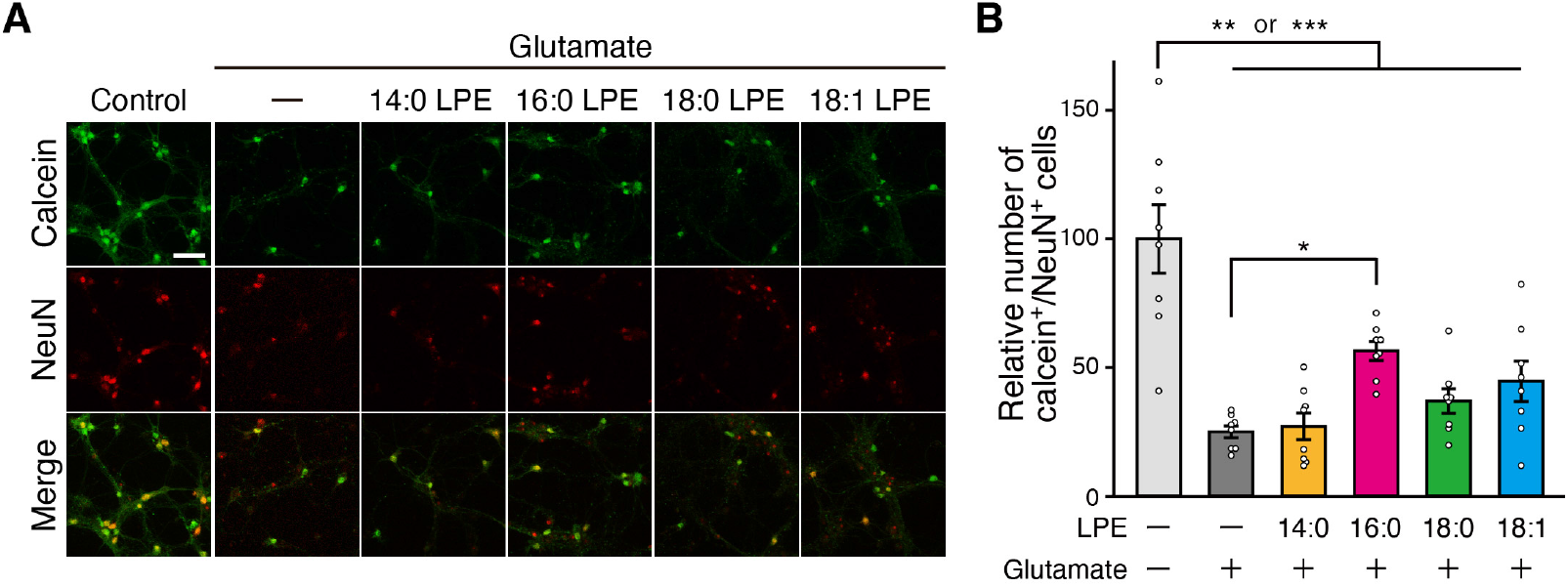
Differential effects of various LPE species on glutamate-induced neurotoxicity in cultured spinal commissural neurons. (A) Effects of LPE species on glutamate-induced neurotoxicity. Cultures were treated with 5 μM glutamate in the presence or absence of 10 μM 14:0 LPE, 16:0 LPE, 18:0 LPE, or 18:1 LPE at DIV 15. After 24 h, cells were stained with calcein and an anti-NeuN antibody. Representative images are shown. Scale bar, 50 µm. (B) Quantification of the number of calcein/NeuN double-positive cells in (A). All values represent mean ± SEM. n = 8 areas from 8 cultures. * p < 0.05, LPE (-) vs 16:0 LPE, **p < 0.01, control vs 16:0 LPE; ***p < 0.001; control vs 14:0 LPE, 18:0 LPE, and 18:1 LPE; one-way ANOVA followed by post hoc Tukey’s test.

### 16:0 LPE reduces apoptotic cell death after SCI

After SCI, extracellular glutamate rapidly increases to toxic levels, leading to excitotoxic neuronal apoptosis. Thus, we examined whether LPE exerts a neuroprotective effect against SCI-induced neuronal cell death in the acute phase of SCI. Among LPE molecular species, 16:0 LPE, which exhibited protective effects against glutamate-induced toxicity in vitro in cultured spinal commissural neurons (Fig. 2) was further evaluated in vivo in SCI model mice. Mice with SCI received intrathecal injections of 16:0 LPE or vehicle immediately after injury (day 0) and again at 1 day post-injury (day 1). At day 2, tissue sections including the spinal cord surrounding the lesion site were prepared and subjected to TUNEL analysis (Fig. 3A and 3B). In the control vehicle-treated mice, numerous TUNEL-positive cells were observed around the SCI region. In contrast, in 16:0 LPE-treated mice, the number of TUNEL-positive cells was significantly reduced to approximately half compared to vehicle-treated mice (Fig. 3C and D). These results suggest that 16:0 LPE also exerts a protective effect on spinal cord neurons against glutamate toxicity-induced cell death in vivo.

**Figure 3.**
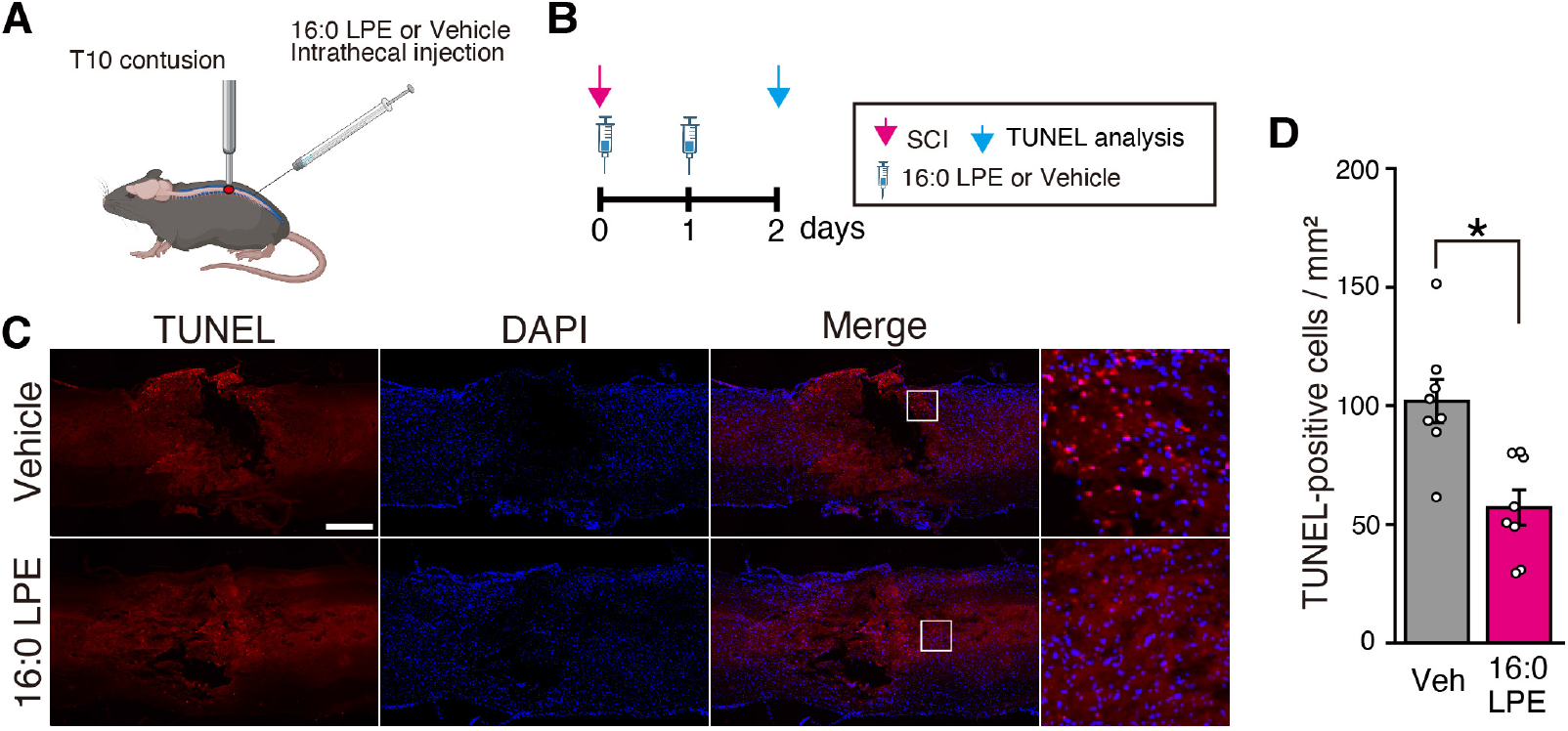
Effect of 16:0 LPE on SCI-induced cell death. (A) Schematic representation of the experimental design used to evaluate the effects of 16:0 LPE on SCI. Contusive SCI was induced at the T10 vertebral level using an impactor device. 16:0 LPE or vehicle is administered intrathecally. (B) Experimental timeline for the TUNEL assay. SCI is induced on day 0, and 16:0 LPE or control vehicle is administered twice, immediately following the SCI on day 0 and once more on day 1. TUNEL analysis is subsequently performed on day 2. (C) Spinal cord sections, including the lesion area, were stained with TUNEL and DAPI. Representative images of TUNEL histochemistry are shown. Boxed regions in the merged panels are shown at higher magnification in the panels on the right. Scale bar, 500 µm. (D) Quantification of TUNEL-positive cells shown in (C). All values presented as mean ± SEM. n = 8 mice per group. *p < 0.05; Welch’s t-test. Veh, vehicle.

### 16:0 LPE reduces lesion expansion following SCI

Next, we performed histological analyses to assess the effect of 16:0 LPE on chronic pathology after SCI. Mice with SCI were administered intrathecal injections of 16:0 LPE or vehicle for a total of three administrations. At 56 days post-injury, spinal cord sections were examined to evaluate the effect of 16:0 LPE on chronic SCI pathology (Fig. 4A). Immunohistochemical analysis for GFAP, an astrocytic marker, and CD11b, a marker for activated microglia and infiltrating macrophages, revealed strong GFAP staining around the injury site, whereas robust CD11b staining was predominantly localized within the GFAP-defined region, in both vehicle-treated and 16:0 LPE-treated mice (Fig. 4B). The region containing CD11b-positive inflammatory cells and delineated by GFAP-positive reactive astrocytes was quantified (Fig.4B–D). In 16:0 LPE-treated mice, the lesion volume defined by this region was significantly smaller than that in vehicle-treated mice (Fig. 4E). Within ± 1.5 mm of the lesion center, total GFAP and CD11b signal intensities showed a trend toward reduction in 16:0 LPE-treated mice compared with vehicle-treated mice, although the differences did not reach statistical significance (Fig. 4F and 4G). These results suggest that 16:0 LPE reduces pathological progression following SCI.

**Figure 4.**
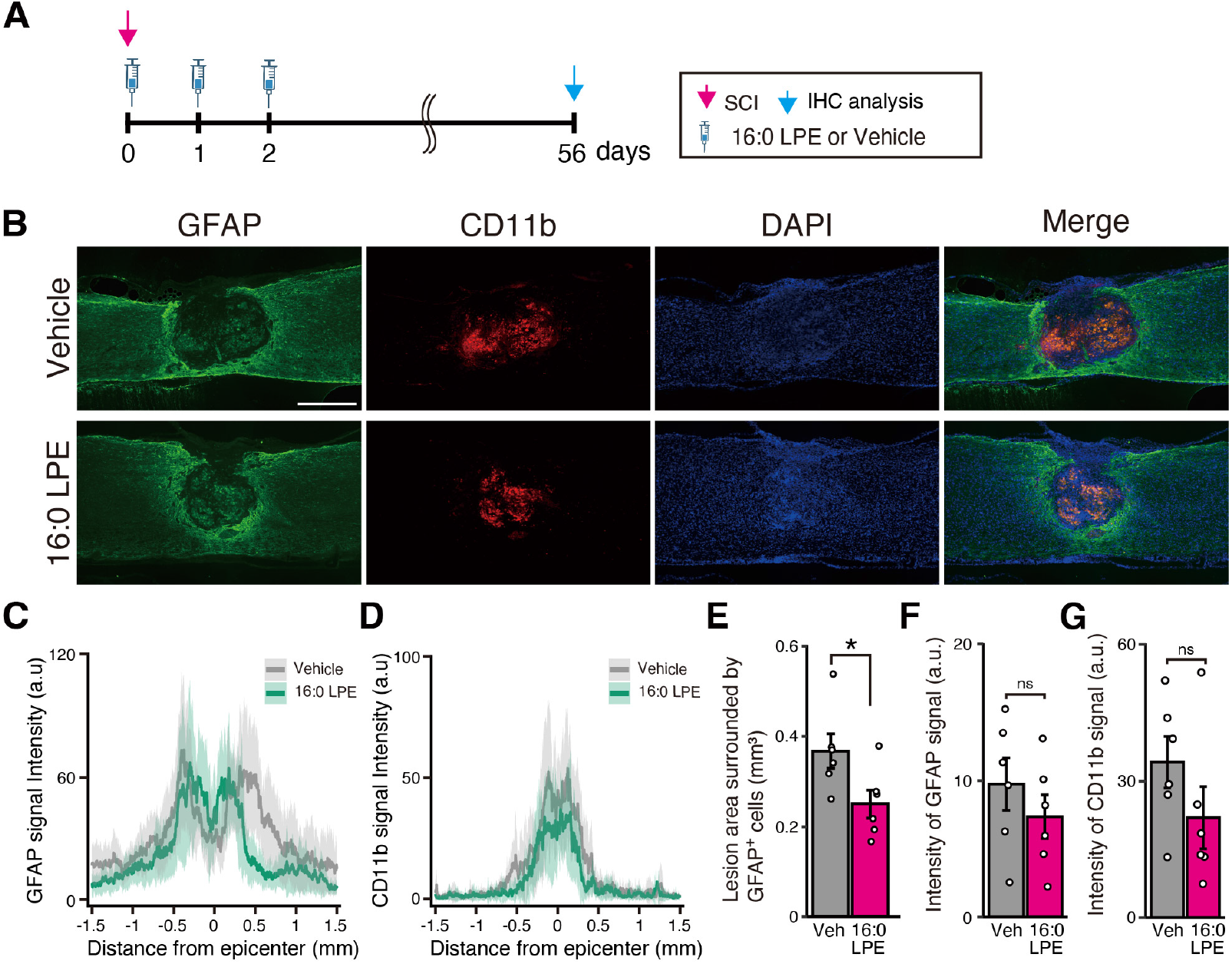
Effect of 16:0 LPE on spinal cord lesion following SCI. (A) Experimental timeline for immunohistochemical analysis of 16:0 LPE effect on SCI-induced lesion. SCI is induced on day 0, and 16:0 LPE or control vehicle is administered three times: immediately after SCI on day 0, on day 1, and on day 2. Immunohistochemical analysis is subsequently performed on day 56. (B) Spinal cord sections, including the lesion area, were immunostained with antibodies against GFAP and CD11b. Representative images are shown. Scale bar, 500 µm. (C) Profile of GFAP signal intensity along the rostro–caudal axis. Signal intensity was measured within ±1.5 mm of the lesion epicenter. (D) Profile of CD11b signal intensity along the rostro–caudal axis. Signal intensity was measured within ±1.5 mm of the lesion epicenter. (E) Quantification of lesion volume. Lesion area, defined by the boundary of GFAP signal, was quantified, and lesion volume was subsequently estimated using five serial sections, including the representative images shown in (B). (E) Quantification of GFAP signal intensity in (B). Measurements were performed within ±1.5 mm of the lesion epicenter. (G) Quantification of CD11b signal intensity in (B). Measurements were performed within ±1.5 mm of the lesion epicenter. All values presented as mean ± SEM. n = 6 mice per group. ns; Welch’s t-test. IHC, immunohistochemistry; Veh, vehicle; ns, not significant.

### 16:0 LPE increases serotonergic fiber innervation following SCI

Serotonergic innervation from the raphe nuclei to the spinal cord has often been associated with improvement in locomotor function. To examine the effect of 16:0 LPE on serotonergic fiber innervation following SCI, spinal cord sections were immunostained for 5-HT at 56 days post-injury (Fig. 4A). In vehicle-treated mice, a weak 5-HT signal was observed on the ventral side of the spinal cord. In contrast, 16:0 LPE-administrated mice exhibited stronger 5-HT signals compared to vehicle-treated controls (Fig. 5A). A rostro–caudal intensity profile demonstrated that the signal increased from the epicenter to 3.0 mm caudal in midsagittal sections (Fig. 5B). Quantitative analysis further revealed that the total 5-HT signal intensity was significantly higher in 16:0 LPE-treated mice compared with vehicle-treated controls (Fig. 5C). Collectively, these results suggested that administration of 16:0 LPE is associated with increased serotonergic innervation following SCI.

**Figure 5.**
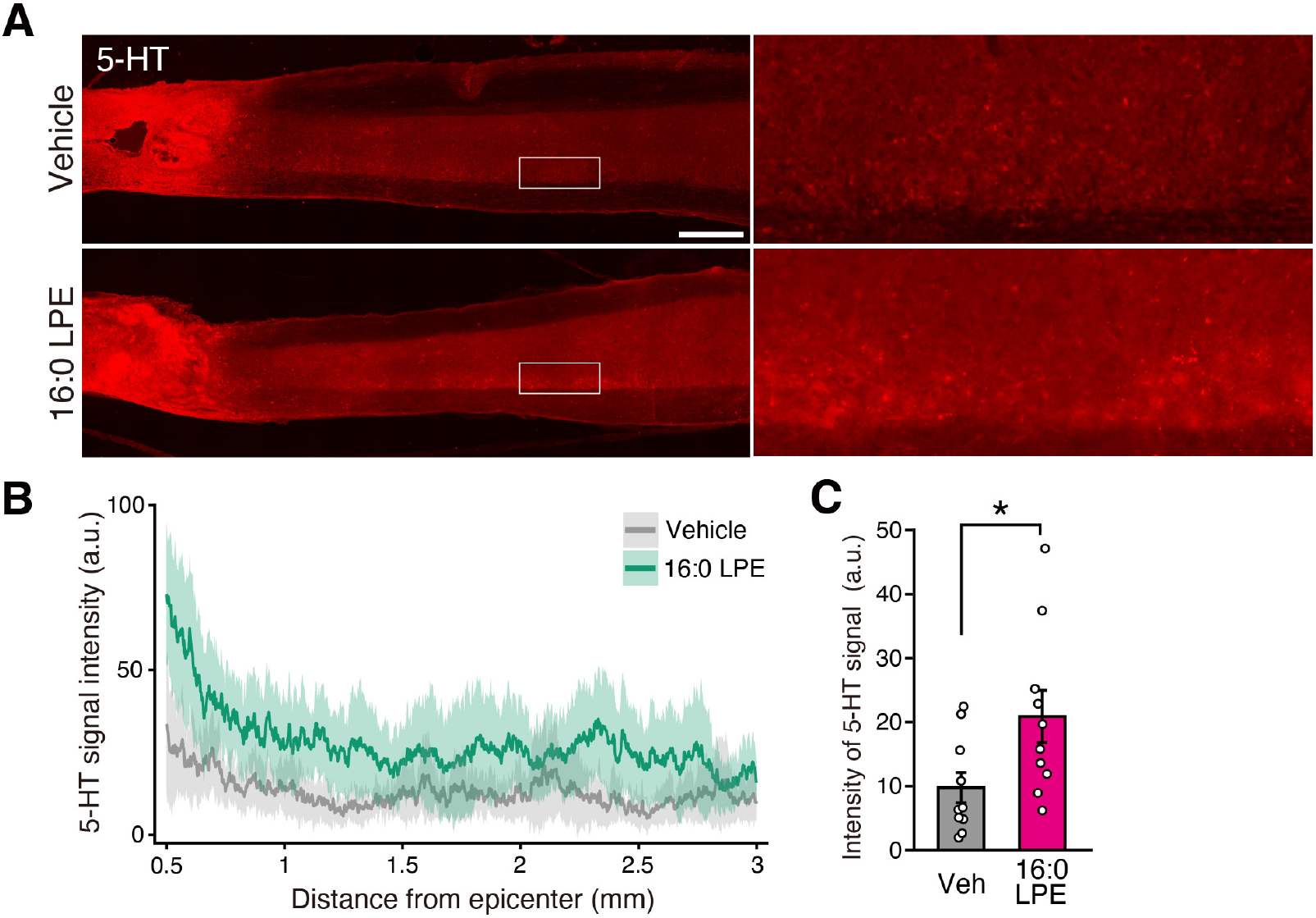
Effect of 16:0 LPE on serotonergic innervation in the spinal cord following SCI. (A) Spinal cord sections were immunostained with antibodies against 5-HT. SCI was induced on day 0, followed by the administration of 16:0 LPE or a vehicle from days 0 to 2. Immunohistochemical analysis was performed on day 56. Representative images are shown. Boxed regions in the merged panels are shown at higher magnification in the panels on the right. Scale bar, 500 µm. (B) Profile of 5-HT signal intensity along the rostro–caudal axis. Signal intensity was measured from 0.5–3.0 mm caudal to the epicenter. (C) Quantification of 5-HT signal intensity in (A). Signal intensity was measured 0.5–3.0 mm caudal to the epicenter. All values presented as mean ± SEM. n = 10 mice per group. *p < 0.05; Welch’s t-test. Veh, vehicle.

### 16:0 LPE promotes locomotor functional recovery following SCI

Finally, we examined whether 16:0 LPE contributes to recovery of locomotor function after SCI. Mice with SCI received intrathecal injections of either 16:0 LPE or vehicle, administered three times in total. Locomotor ability was then monitored weekly for eight weeks (Fig. 6A). Motor function was evaluated using the Basso Mouse Scale (BMS), which assesses hind-limb function. Both vehicle-treated mice and 16:0 LPE–treated mice showed a gradual increase in BMS scores from week 1 to week 8 (Fig. 6B). However, the BMS scores of 16:0 LPE–treated mice were significantly higher compared to those of vehicle-treated mice. These results suggested that 16:0 LPE facilitates functional recovery of locomotion after SCI.

**Figure 6.**
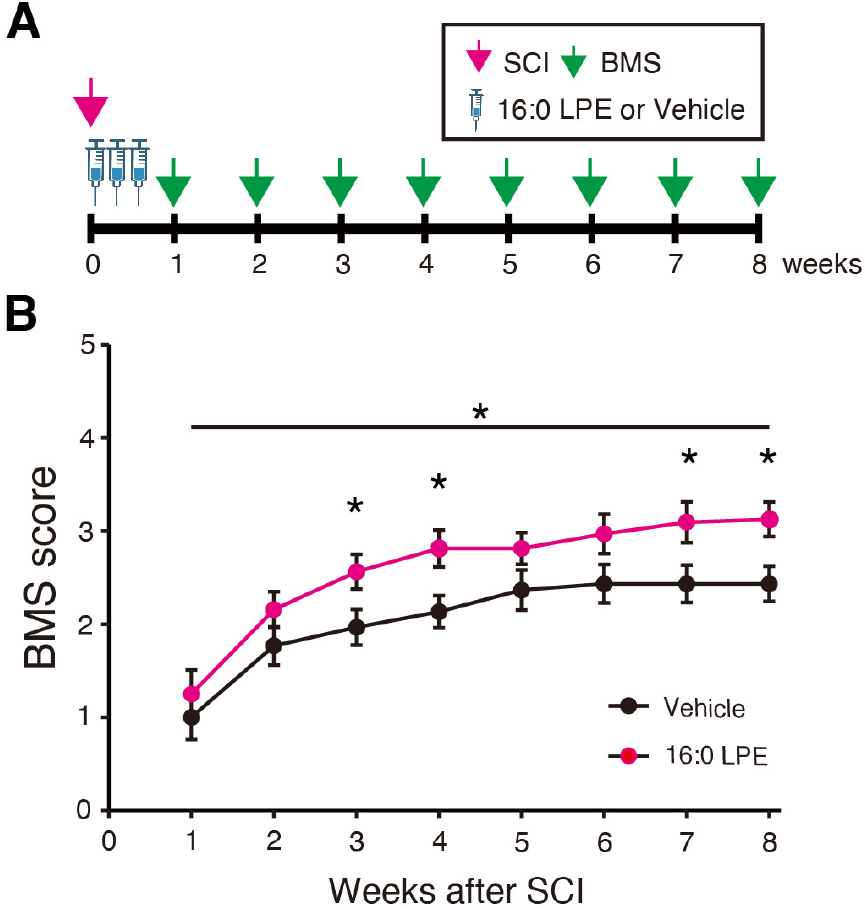
Effect of 16:0 LPE on locomotor recovery following SCI. (A) Experimental timeline. SCI was induced on day 0, and 16:0 LPE or vehicle was administered three times: immediately after SCI (day 0), on day 1, and on day 2. Locomotor function was assessed weekly for eight weeks. (B) Locomotor recovery assessed using the BMS. All values presented as mean ± SEM. n = 15 for vehicle; n = 16 for 16:0 LPE. *p < 0.05; two-way repeated-measures ANOVA followed by post hoc Welch’s t-test.

## Discussion

The present study yielded four principal findings. First, LPE exerted neuroprotective effects against glutamate toxicity, which varied among molecular species, with 16:0 LPE showing the most pronounced effect in spinal cord neurons. Second, histological analysis revealed a significant reduction in apoptotic cells at two days post-injury in the LPE-treated group compared with the vehicle group. Third, at eight weeks post-injury, lesion volume was reduced and serotonergic innervation was enhanced in LPE-treated mice. Finally, intrathecal administration of 16:0 LPE resulted in significantly improved locomotor recovery over the eight-week observation period.

A variety of LPE species, differing in fatty acyl chain length and saturation, have been identified in mammalian tissues (Van Blitterswijk et al., 1982). Although no definitive receptor has been assigned to LPE, increasing evidence suggests that distinct G protein–coupled receptors (GPCRs), expressed in a cell type–specific manner, may mediate its effects (Hisano et al., 2021a; Hisano et al., 2021b; Park et al., 2007; Park et al., 2014). Experimental studies indicate that 14:0, 16:0, and 18:1 LPE can activate MAPK/ERK1/2 signaling in PC12 cells (Nishina et al., 2006), while 18:1 LPE activates ERK1/2 in MDA-MB-231 breast cancer cells (Park et al., 2014). More recently, our group reported that 18:0 LPE acts through a Gi/Go-coupled GPCR–PLC pathway (Hisano et al., 2021a), that 18:1 LPE promotes neurite outgrowth via a Gq/11–PLC–PKC–MAPK cascade (Hisano et al., 2021b), and that 16:0 LPE signals through a Gq/11-coupled GPCR linked to PLC activation (Hisano et al., 2021a). Brief ERK1/2 activation by neurotrophic factors has been shown to reduce glutamate toxicity (Subramaniam and Unsicker, 2010), which may underlie the observed neuroprotective effects of LPE. Further studies are required to identify the receptors responsible for mediating LPE signaling.

Following SCI, extracellular glutamate levels rapidly increase to excitotoxic concentrations and remain elevated for days to weeks (Panter et al., 1990; Park et al., 2004; Xu et al., 2004). Such elevations contribute to neuronal and glial apoptosis (Ahuja et al., 2017). Our results indicate that 16:0 LPE reduces neuronal apoptosis in the acute phase and limits lesion expansion in the chronic phase. These findings are consistent with the hypothesis that LPE mitigates excitotoxicity and secondary injury. LPE may also exert indirect benefits by modulating neuroinflammation. Previous studies have reported that LPE and other lysophospholipids display anti-inflammatory and antioxidant properties in activated microglial cells and peripheral models of inflammation (Hung et al., 2011; Tsukahara et al., 2021; Park and Im, 2021). Therefore, the beneficial effects of LPE in SCI may involve both neuronal and non-neuronal mechanisms.

The observed increase in serotonergic innervation following LPE administration may contribute to functional recovery. Whether this increase reflects preservation of pre-existing fibers or promotion of axonal sprouting remains unclear. Since regeneration in the central nervous system generally requires weeks to months (Nagappan et al., 2020; Tsujioka and Yamashita, 2021; Varadarajan et al., 2022), preservation of serotonergic projections is the more parsimonious explanation. However, our in vitro findings indicate that LPE, particularly 16:0 and 18:1 species, can promote neurite extension, raising the possibility that LPE also supports regenerative growth after SCI.

In summary, intrathecal administration of 16:0 LPE reduced neuronal apoptosis, limited lesion progression, enhanced serotonergic innervation, and improved locomotor function following SCI. These findings suggest that 16:0 LPE is a promising candidate for neuroprotective therapy and provide new insight into the actions of lysophospholipids in the central nervous system.

## Acknowledgements

We thank the staffs of the Animal Facility of Nihon Pharmaceutical University. This work was the result of using the research equipment shared in the MEXT Project for promoting public utilization of advanced research infrastructure (Program for supporting the construction of core facilities; grant number: JPMXS0441000021). Some figures were created with BioRender.com.

## Authors contributions

T.M. and T.K. performed behavioral experiments. T.Y. and S.K. performed cell culture experiment and analyzed the data from the behavioral experiments. T.M. and T.U. wrote the manuscript, which was edited by E.K., T.K., Y.M., N.S., and J.T. T.M. and T.U. designed the study, and T.U. supervised the study.

## Data availability statement

The data generated in this study are available from the corresponding author upon reasonable request.

